# The ATP-dependent protease ClpYQ degrades cell division proteins DivIVA and Mbl in *Bacillus subtilis*

**DOI:** 10.1101/2024.10.15.618472

**Authors:** Taylor D. Lanosky, Michael G. Darnowski, Jordan T. Brazeau-Henrie, Puneet Labana, Christopher N. Boddy

## Abstract

ATP-dependent proteases play key roles in bacterial protein quality control and regulation of cellular processes. ClpYQ and ClpXP are ATP-dependent proteases in the Gram-positive bacteria *Bacillus subtilis*. To date, no substrate proteins of *B. subtilis* ClpYQ have been characterized. The protease component encoded by *clpQ* is synthetically lethal with *clpP* and the two genes are non-essential individually, suggesting potentially redundant roles for ClpYQ and ClpXP. Previous quantitative proteomic data predicted that *B. subtilis* proteins DivIVA and Mbl, components of the divisome and elongasome respectively, are potential substrates of ClpYQ. The role of DivIVA and Mbl in cell division and elongation suggests a significant role of ClpYQ in regulating cell division through targeted degradation of key divisome and elongasome proteins. Here we confirm that DivIVA and Mbl are degraded by ClpYQ both *in vitro* and *in vivo*, and thus identify the first two substrates of ClpYQ in *B. subtilis*.

## Introduction

AAA+ proteases (ClpXP, ClpAP, ClpCP, ClpYQ, Lon, FtsH, PAN/20S, and the 26S proteasome) play key roles in bacterial protein quality control and regulation of cellular processes [1,2]. All AAA+ family proteolytic machines contain a hexameric ring of AAA+ ATPases (chaperones) and a compartmental protease [1]. Exposed segments of substrate proteins (called degradation tags or degrons) are bound to the axial pore of the AAA+ ring either directly or via adaptor proteins. ATP binding and hydrolysis powers conformational changes in the ring that creates an unfolding force and translocates the unfolded substrate into the peptidase compartment for degradation [1,3]. Since degradation is irreversible, highly specific substrate recognition is essential.

In the Gram-positive bacterium *Bacillus subtilis*, there are six well-characterized AAA+ proteases: ClpCP, ClpEP, ClpXP, LonA, LonB, and FtsH [3]. Substrates of ClpXP have been successfully identified through proteomic analyses. Initially this involved comparison of the proteomes of wildtype strains to those of *clpX* and/or *clpP* mutant strains using two-dimensional polyacrylamide gel electrophoresis [4,5]. More recently, trapping of endogenous substrates within inactivated ClpP *in vivo* followed by purification of the trapped protein complex and subsequent identification by mass spectrometry has also been used [6,7]. Substrates identified from these types of proteomic experiments must still be validated *in vitro* and *in vivo* to confirm their roles as client proteins. *In vitro* characterization typically involves degradation of recombinant purified putative substrates by recombinant purified ClpXP [6,8,9]. For *in vivo* validation, a common strategy is inhibition of total protein synthesis in cultures of wildtype and *ΔclpX* strains followed by quantification of client proteins by western immunoblotting [6,9].

ClpYQ, also known as HslUV or CodWX, is another ATP-dependent protease in *B. subtilis*. It is composed of a 50 kDa hexamer of the ATPase ClpY bound to a 19 kDa hexamer of the protease ClpQ [10–12]. Native protein substrates are recognized and unfolded by ClpY, and the unfolded substrate is degraded by ClpQ through an N-terminal active site serine [10,13]. The biological function and regulation of ClpYQ in *B. subtilis* are poorly understood. One study demonstrated that a *ΔclpYQ* mutant formed early and robust biofilms, suggesting that ClpYQ regulates biofilm formation [14]. An absence of ClpYQ has also been shown to impair swarming motility [14,15]. In *Staphylococcus aureus*, another Gram-positive bacterium, ClpYQ has been suggested to play a minor role in virulence [16]. The protease component encoded by *clpQ* is synthetically lethal with *clpP*, indicating essential and potentially redundant roles for ClpYQ and ClpXP [17]. Recently, we showed that the antibiotic armeniaspirol inhibits ClpYQ and ClpXP *in vitro* and in *B. subtilis*, leading to a dysregulation of cell division [17,18].

Unlike ClpXP, there are no confirmed substrate proteins of ClpYQ. Through global proteomic analysis using isobaric tags for relative and absolute quantification (iTRAQ), one study found that various proteins involved in motility, chemotaxis, and the production of γ-PGA were less abundant in a *B. subtilis ΔclpYQ* mutant compared to the wildtype strain [14]. Another study from our lab used dimethyl isotopic labeling quantitative proteomics to show that several components of the divisome and elongasome such as DivIVA and Mbl increased in abundance in a *B. subtilis ΔclpQ* mutant compared to the wildtype strain (Table S1) [17]. Based on our data, we set out to determine if DivIVA and Mbl are substrates of ClpYQ.

The divisome is a contractile protein ring that promotes invagination of the cytoplasmic membrane and cell wall, and mediates formation of the cell division septum [19–21]. Divisome assembly is directed by FtsZ, which polymerizes at the mid-cell to form a Z-ring upon which all other divisome-related proteins are scaffolded, guiding septum synthesis [22,23]. It assembles concomitantly with FtsA, ZapA, and EzrA into the early divisome. DivIVA and numerous extracytoplasmic proteins are then recruited to form the late divisome [24]. Due to its high affinity for negative membrane curvature, DivIVA localizes to polar regions and nascent division sites where it complexes with MinC and MinD to inhibit FtsZ polymerization [25–29]. DivIVA has also been suggested to interact with Maf to inhibit cell division during competency [30]. Furthermore, DivIVA helps coordinate chromosome segregation [31,32]. Overexpression of DivIVA in *B. subtilis* is lethal, indicating that dysregulation of the divisome can be catastrophic [33].

The elongasome directs peptidoglycan hydrolysis and synthesis along the long axis of the cell, enabling cylindrical growth [34]. In *B. subtilis*, it is guided by the redundant action of three MreB isoforms: MreB, Mbl, and MreBH [35–37]. Under normal growth conditions, MreB and Mbl are essential [38]. MreBH is only essential under certain adverse conditions [35,36]. In *mbl* mutants, the Z-ring of the divisome cannot properly close upon itself and instead forms distorted structures [39]. Cell division is inhibited by MreB in *B. subtilis*, and overexpression of MreB or MreBH is lethal [36,38,40].

Using both *in vitro* and *in vivo* degradation assays, we identify DivIVA and Mbl as the first two substrates of the AAA+ protease ClpYQ. Given that DivIVA and Mbl are key components of the divisome and elongasome, this study suggests a critical role for ClpYQ in regulating cell division. These findings also suggest that Mbl may be a substrate of LonA and that DivIVA and Mbl are substrates of ClpXP, highlighting the redundancy of ATP-mediated proteolysis of substrates that play an essential role in cell division and elongation. These discoveries help illustrate the role of ClpYQ, an ATP-dependent protease that was previously uncharacterized.

## Results

### C-terminally His-tagged Mbl and DivIVA are degraded *in vitro*

To evaluate if Mbl and DivIVA are degraded by ClpYQ *in vitro*, degradation assays were conducted with recombinant purified protein to screen for a change in abundance in the presence of ATP relative to a no-ATP control. Trigger Factor (TF) and EzrA were selected as negative controls since previous quantitative proteomics data predicted that they were not ClpYQ substrates (Table S1) [17]. The assay was conducted using recombinant purified proteins with both N- and C-terminal His tags since the degron, which is required for substrate recognition by ClpYQ, could be on either terminus. Of the eight proteins, only the N-terminally His-tagged DivIVA construct could not be successfully expressed and purified. Digests were analyzed by SDS-PAGE and quantification of the protein bands showed statistically significant degradation of C-terminally His-tagged Mbl and DivIVA compared to non-client proteins (Fig. 1A). A ClpYQ inhibitor, Cl-armeniaspirol, was added to ensure that C-terminally His-tagged Mbl and DivIVA were not degraded in a ClpYQ-independent manner [18]. There was a statistically significant decrease in the degradation of client proteins in the presence of Cl-armeniaspirol compared to in its absence, and there was not a statistically significant difference in their abundance in the presence of the inhibitor compared to the abundance of non-client proteins (Fig. S1). The N-terminally His-tagged Mbl was not degraded by ClpYQ (Fig. 1B). These results are consistent with DivIVA and Mbl being ClpYQ substrates and suggest that substrate recognition occurs at these targets’ N-termini.

**Figure 1.**
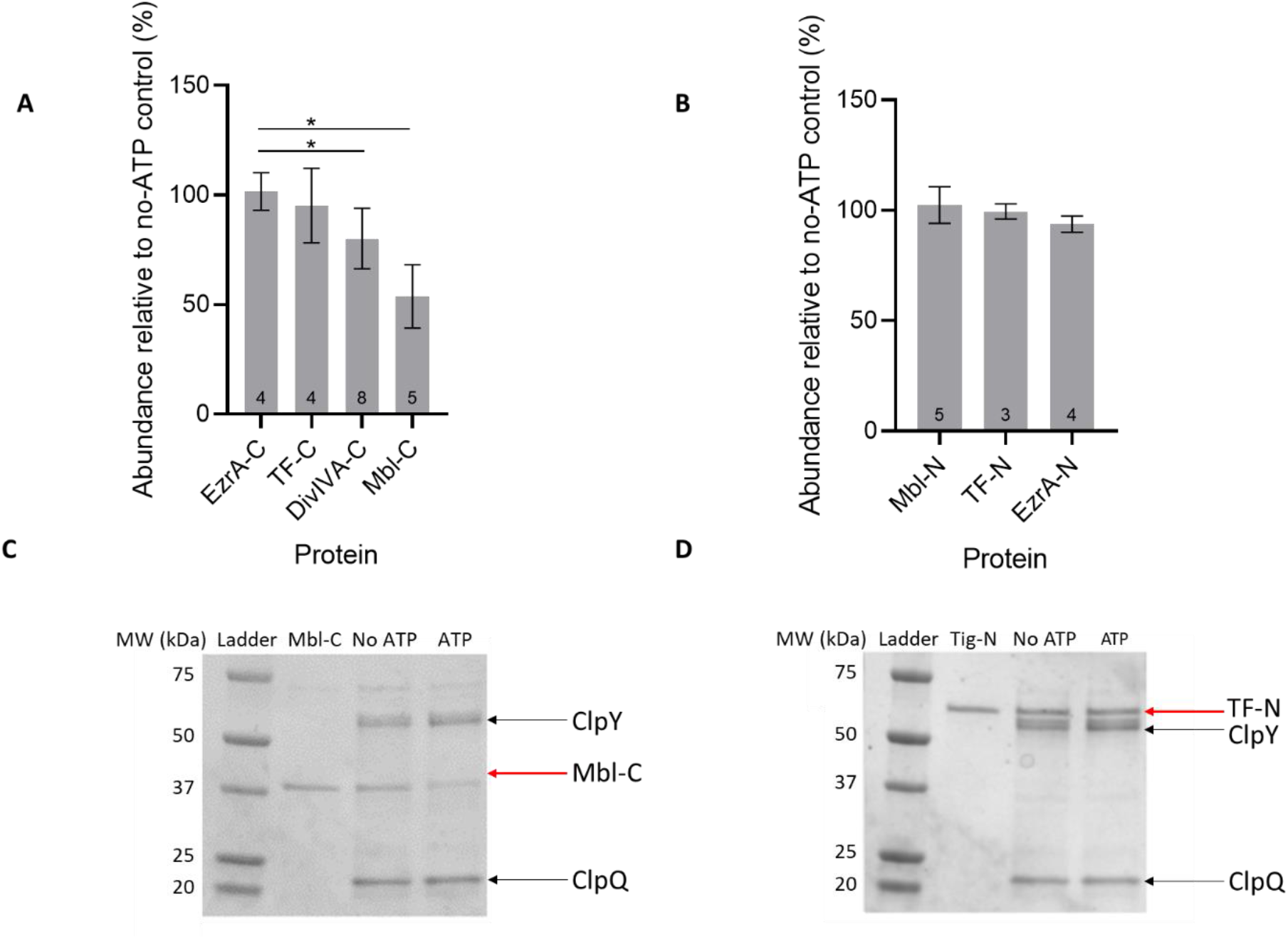
*In vitro* degradation of C-terminally His-tagged Mbl and DivIVA by ClpYQ. 2.5 µM of recombinant purified Mbl and DivIVA were incubated with 5 µM of recombinant purified ClpY and ClpQ for 24 h at 37 °C. TF and EzrA were used as negative controls. The terminus onto which the His tag was fused is indicated by the C (***A***) or N (***B***) suffix on the protein names. Digests were analyzed by SDS-PAGE. Each band was normalized to a loading control before quantifying the amount of protein remaining in the reaction mixture relative to a no-ATP control. Error bars represent the standard deviations from n=3-8 independent replicates. All groups were compared to each other using a one-way ANOVA with a Tukey-Kramer HSD variation to ensure a statistically significant difference in remaining protein abundance between client and non-client proteins (Table S5). Asterisks (^*^) represent statistical significance (p≤ 0.05). The p-value between EzrA-C and DivIVA-C was 0.012, while the p-value between EzrA-C and Mbl-C was 8.83 × 10^−7^. ***C***. Representative SDS-PAGE gel of a ClpYQ client protein (Mbl-C). Decreased protein abundance is observed in the presence of ATP due to ClpYQ-mediated degradation. Both here and in ***D***, the first lane after the ladder (labeled by the protein name) represents the loading control. ***D***. Representative SDS-PAGE gel of a non-client protein of ClpYQ (TF-N). No change in protein abundance is observed in the presence of ATP, indicating no ClpYQ-mediated degradation.

### Mbl and DivIVA are degraded *in vivo*

We designed an assay to evaluate if DivIVA and Mbl are ClpYQ substrates *in vivo*. Because *clpP* and *clpQ* are synthetically lethal, we suspected ClpYQ and ClpXP may be able to complement each other. This is supported by the observation that DivIVA is a substrate of ClpXP in *Streptococcus mutans* [41]. Thus to evaluate ClpYQ activity independent of ClpXP, we used a *ΔclpP* strain, eliminating all ClpXP activity. This strain was transformed with expression vectors encoding C-terminally His-tagged Mbl or DivIVA under the control of an IPTG-inducible promoter. Treatment of these transformants with or without the ClpYQ inhibitor Cl-armeniaspirol enabled evaluation of *in vivo* ClpYQ proteolysis of the client proteins. Expression of DivIVA and Mbl was induced for 6 h before inhibiting total protein translation using spectinomycin at its MIC (32 µg/mL and 64 µg/mL for the strain transformed with Mbl and DivIVA, respectively). At the same time protein translation was inhibited, half the MIC of Cl-armeniaspirol was added to antibiotic-treated cells. The MICs of Cl-armeniaspirol in the *ΔclpP* strain transformed with Mbl and DivIVA were 2 µg/mL and 4 µg/mL, respectively. Whole cell lysates obtained at different time points after the inhibition of translation were separated by SDS-PAGE and proteins were visualized by western blot analysis. Quantitative analysis of protein bands showed a statistically significant decrease in abundance of DivIVA and Mbl 90 min after the inhibition of protein synthesis in cells without Cl-armeniaspirol compared to cells with Cl-armeniaspirol (Fig. 2A,C). These results are consistent with our *in vitro* results. Significantly more degradation of Mbl than DivIVA was observed in both the presence and absence of Cl-armeniaspirol (Table S6).

**Figure 2.**
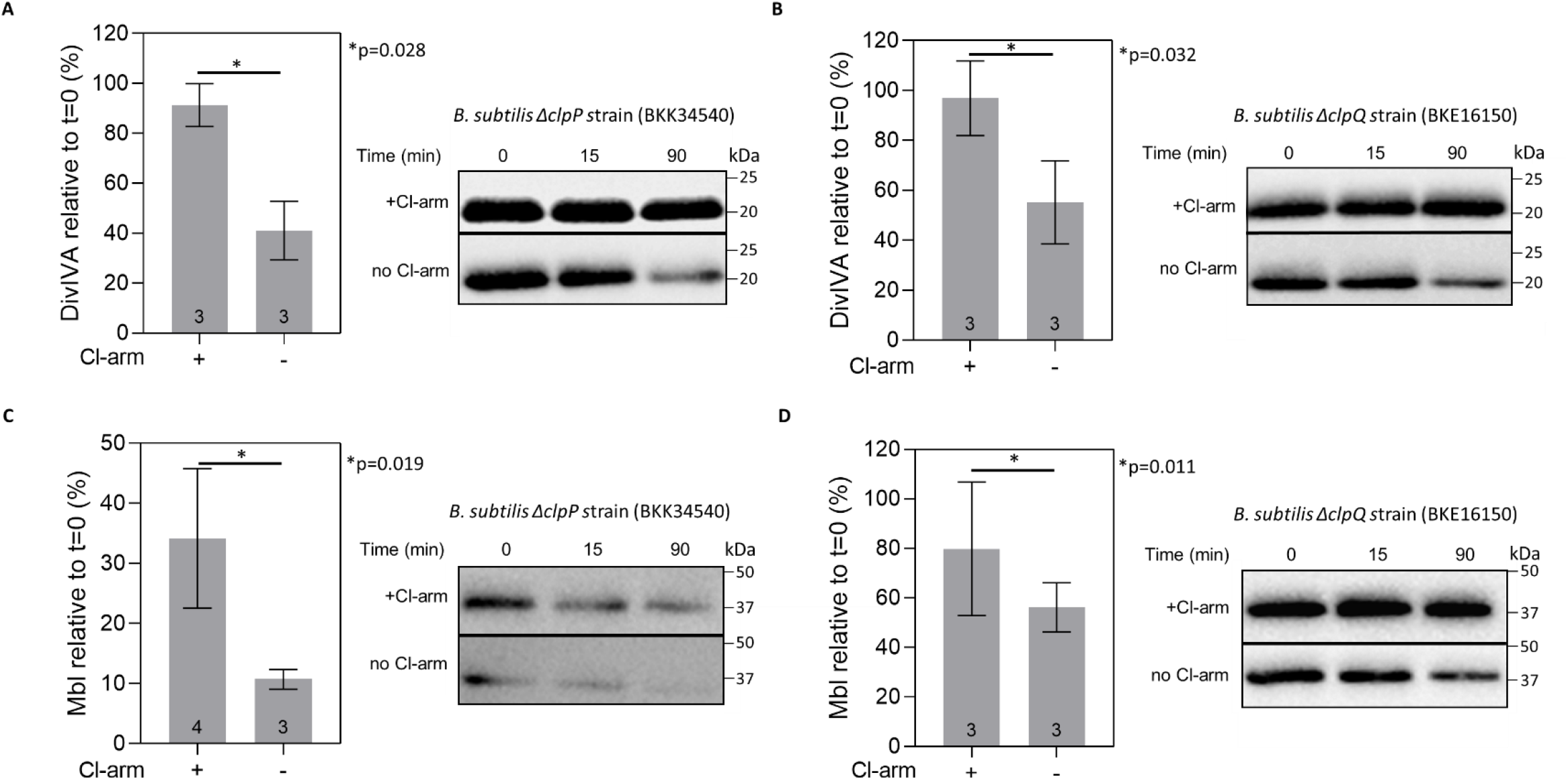
Cellular degradation of DivIVA and Mbl. ***A***. Western blot analysis of degradation of overexpressed DivIVA as a function of time after inhibition of protein synthesis in *ΔclpP* cells with (+) or without (-) Cl-armeniaspirol. Cl-ameniaspirol was added to inhibit ClpYQ in control cells. The bar graph on the left shows quantification of the protein bands at t=90 min relative to t=0 min. Reactions separated by solid lines are from different gels. Error bars here and in ***B-C*** represent the standard deviations from n=3-4 individual replicates. Student’s two-tailed t-test was used to compare results from each strain. The time point taken directly after inhibition of protein synthesis refers to t=0 min. ***B***. Same as in ***A*** but in *ΔclpQ* cells and Cl-armeniaspirol was added to inhibit ClpXP in control cells. ***C***. Same as in ***A*** but using overexpressed Mbl. ***D***. Same as in ***A*** but using overexpressed Mbl in *ΔclpQ* cells and Cl-armeniaspirol was added to inhibit ClpXP in control cells.

To evaluate the hypothesis that ClpXP and ClpYQ can complement each other, we examined the ability of ClpXP to degrade Mbl and DivIVA *in vivo*. A *ΔclpQ* strain was transformed with an expression vector encoding C-terminally His-tagged Mbl or DivIVA, respectively. In the *ΔclpQ* mutant, ClpYQ cannot assemble to mediate degradation of its client proteins; this allows for the exclusive analysis of ClpXP-mediated degradation of Mbl and DivIVA. In these assays, Cl-armeniaspirol was added to control cells to inhibit ClpXP. The MICs of spectinomycin for the *ΔclpQ* strain transformed with Mbl and DivIVA were 32 µg/mL and 128 µg/mL, respectively, and of Cl-armeniaspirol were both 2 µg/mL. Data showed a statistically significant decrease in abundance of DivIVA and Mbl 90 min after the inhibition of protein synthesis in cells without Cl-armeniaspirol compared to cells with Cl-armeniaspirol (Fig. 3B,D). These data strongly support the hypothesis that Mbl and DivIVA are also substrates of ClpXP. Mbl and DivIVA were not confirmed as ClpXP substrates in *in vitro* degradation assays because recombinant purified *B. subtilis* ClpXP is non-functional [17].

## Discussion

Through a combination of *in vitro* and *in vivo* degradation assays, DivIVA and Mbl have been confirmed as the first substrates of the *B. subtilis* AAA+ protease ClpYQ. These results support previous quantitative proteomics-based predictions [17].

We have shown that ClpYQ-mediated *in vitro* degradation is observed only for C-terminally His-tagged DivIVA and Mbl. The successful degradation of DivIVA and Mbl indicates that adaptor proteins are not required for these client proteins and that substrate recognition occurs at the N-termini of these client proteins. Previous quantitative proteomics data identified numerous proteins in addition to Mbl and DivIVA that increased in abundance in a *B. subtilis ΔclpQ* strain [17]. However, alignment of the N-termini of these putative ClpYQ clients (including Mbl and DivIVA) showed no conserved sequence element. Furthermore, analysis with Multiple Em for Motif Elicitation (MEME) [42] also did not identify any conserved sequence elements. The nature of degrons have not been analyzed in depth for all AAA+ protease substrates. In some cases they are well defined. For example, it is known that five classes of conserved degrons exist for *E. coli* ClpXP: two located at the C-terminus and three at the N-terminus of proteins [6]. In contrast some recognition elements are very broad. For example, a broad set of hydrophobic sequences that are preferably rich in aromatic residues mediates substrate recognition by *E. coli* AAA+ protease Lon [43–46]. Thus, a lack of conserved degron sequences for ClpYQ is not atypical.

In designing our *in vivo* assay to evaluate ClpYQ degradation of DivIVA and Mbl, we hypothesized that these client proteins could also be substrates of ClpXP. This hypothesis is supported by observations that *clpQ* is synthetically lethal with *clpP*, suggesting redundant roles for ClpYQ and ClpXP [17]. Moreover, DivIVA is a proposed substrate of ClpXP in the Gram-positive bacteria *Streptococcus mutans* [41]. Furthermore, quantitative proteomics of a *B. subtilis ΔclpP* mutant versus the wild-type strain identified an increased abundance of DivIVA in two out of three replicates and a 16-fold increase in abundance of Mbl with a p-value of 0.052 from a two-tailed Students t-test (Table S1) [17]. While not significant in the two-tailed t-test, this result reaches statistical significance in the one-tailed t-test which may be more appropriate since the hypothesis predicts an increase in Mbl levels in the *ΔclpP* mutant [17]. Compensatory activity of proteases is not uncommon. For example, in *B. subtilis* itself, the HtrA and HtrB proteases have been shown to compensate for one another, and it has been suggested that HtrC can compensate for the lack of both HtrA and HtrB [47]. It has also been proposed that LonA and LonB could compensate for one another in *B. subtilis* [48]. In the cyanobacterium *Synechoccus elongatus*, compensatory effects observed between the Clp proteases ClpP1, ClpP2, ClpP3, and/or ClpR are suggested to be important for determination of the circadian period and viability [49,50]. There was thus substantial literature precedent for this hypothesis.

Our *in vivo* degradation assays performed in the *ΔclpP* mutant removed any confounding effect of ClpXP-mediated degradation and confirmed that DivIVA and Mbl are substrates of ClpYQ *in vivo* (Fig. 2A,C). Since substantially more degradation of Mbl than DivIVA was observed in both the presence and absence of Cl-armeniaspirol, it is likely that Mbl is a potential substrate of another protease. Given that the *ΔclpP* strain was used for these assays, the protease responsible for the increased degradation cannot be ClpCP, ClpEP, or ClpXP. Although it could be FtsH or LonB, these proteases are anchored to the cellular membrane [51– 54] and may be less likely to degrade Mbl as a cytosolic target. Thus, we speculate that Mbl may also be a substrate of LonA.

*In vivo* degradation assays of Mbl and DivIVA in *ΔclpQ* mutant cells showed decreased abundance of DivIVA and Mbl compared to Cl-armeniaspirol treatment. In *ΔclpQ* mutants, the effects of ClpYQ-mediated degradation are removed and the decrease in Mbl and DivIVA abundance can likely be attributed to ClpXP-mediated degradation. Thus, these results provide further support for our hypothesis that DivIVA and Mbl are substrates of ClpXP. Slightly more degradation of DivIVA and substantially more degradation of Mbl was mediated by ClpYQ than by ClpXP, respectively, suggesting that DivIVA and Mbl have higher affinities for ClpYQ than ClpXP. Bioinformatic analysis was also performed on the N-terminus of proteins that increased in abundance in a *B. subtilis ΔclpP* strain according to previous quantitative proteomics data [17], and no motifs were identified to act as likely degradation tags for ClpXP.

Substrate redundancy by ClpYQ and ClpXP may be important for controlling cell division. DivIVA is a key component of the divisome and inhibits Z-ring formation, which ultimately inhibits cell division [25–29]. For these reasons, overexpression of DivIVA in *B. subtilis* is lethal [33]. Mbl and its isoforms, MreB and MreBH, guide the elongasome [35–37]. In *B. subtilis*, overexpression of the Mbl isoforms MreB and MreBH are lethal, and cell division is inhibited by MreB [36,38,40]. Given the catastrophic effects that arise upon the overexpression of these proteins, ensuring regulated proteolysis is essential. Unlike genetic tools, Cl-armeniaspirol overcomes ClpYQ/ClpXP redundancy by inhibiting both as has previously been shown [17,18].

In summary, this study identifies DivIVA and Mbl as the first two confirmed substrates of ClpYQ in *B. subtilis*. The importance of these substrates in cell division and elongation suggests that ClpYQ plays an important role in regulating cell division through targeted degradation of key divisome and elongasome proteins. This data further highlights the utility of the ClpYQ inhibitor Cl-armeniaspirol in studying AAA+ proteases. We also provide preliminary data identifying DivIVA and Mbl as potential targets of ClpXP and propose Mbl to be a target of LonA. Taken together, these results help elucidate the role of the previously uncharacterized ATP-dependent ClpYQ: an important protease for the proper function and viability of *B. subtilis*.

## Experimental procedures

### Cloning and protein purification

*B. subtilis clpY, clpQ, divIVA, mbl, tig*, and *ezrA* genes were PCR amplified from *B. subtilis* 168 genomic DNA (Promega Wizard Genomic DNA Purification Kit). Primers used are shown in Table S2. Cloning of *clpQ*, its encoded protein expression and purification were performed as previously explained by Labana and collaborators [17]. The *clpY* gene was cloned into pET28b (N-terminal His tag, KanR). The resulting plasmid (pPL31) was transformed into chemically competent *E. coli* BL21 (DE3) for protein expression. 400 mL LB media was inoculated with 0.5% (v/v) of an overnight pre-culture. The culture was grown at 37°C (200 rpm) and expression was induced with 1 mM isopropyl-1-thio-β-D-galactopyranoside at an optical density (OD_600_) of 0.7. The culture was grown at 16°C (200 rpm) overnight. The cells were pelleted at 4,000 rpm for 20 min and re-suspended in lysis buffer (100 mM Tris, 300 mM NaCl, 1 μg/mL leupeptin, 1 μg/mL pepstatin A, 1 mg/mL lysozyme, 10% glycerol, pH 8.0). Mechanical cell lysis was achieved by three 1 minute cycles of 3 s sonication and 2 s rest on ice followed by a 1 min incubation on ice. The cell debris were pelleted at 10,000 rpm for 60 min and the supernatant was incubated with 800 μL 50% Ni-NTA agarose resin (QIAGEN) for 1 h at 4°C with gentle shaking. The lysate was loaded onto a column and the flow-through was collected. The resin was washed sequentially with 5 mL elution buffer (100 mM Tris, 300 mM NaCl, pH 8.0) containing 0 mM, 20 mM, 100 mM and 250 mM imidazole. Fractions were analyzed by SDS-PAGE (4-20% Mini-PROTEAN TGX Precast Gels; Bio-Rad). ClpY-containing fractions were pooled before the protein was concentrated and elution buffer was exchanged with dialysis buffer (300 mM NaCl, 100 mM Tris, pH 8.0) using a 2kDa Slide-A-Lyzer® Dialysis Cassette (Thermo Scientific). The concentrated protein was stored at -80°C with 30% (w/w) glycerol. All protein concentrations were determined by Bradford assay.

The *mbl, divIVA, tig, and ezrA* genes were cloned into PET21b vectors (C-terminal His tag, AmpR) and pMGXHA vectors (N-terminal His tag, AmpR). The resulting plasmids (pTL01-pTL08) were expressed and purified the same way as pPL31, but protein concentration and buffer exchange were done using a 10 kDa Amicon Ultra-15 centrifugal filter unit (Millipore Sigma). The concentrated protein was stored at -80°C with 30% (w/w) glycerol.

For *in vivo* studies, *mbl* and *divIVA* fused with C-terminal His tags were PCR amplified from pTL01 and pTL02, respectively, and cloned into the pHT01 *B. subtilis* expression vector (inducible σA-dependent promoter, CmR). Primers used are shown in Table S2. The resulting plasmids were pTL12 and pTL13, respectively. A summary of all plasmids constructed in this study is shown in Table S3.

### *In vitro* degradation assays

*In vitro* degradation was evaluated by incubating recombinant purified proteins (2.5 µM) with recombinant purified ClpY and ClpQ (both 5 µM) in reaction buffer for 24 h at 37 °C. Reaction buffer was composed of 0.1 M Tris (pH 8.0), 10 mM MgCl_2_, 5 mM ATP, 1 mM TCEP, and 1 mM EDTA. Approximately 4X the K_I_ of Cl-armeniaspirol for ClpYQ, or 25 µM of Cl-armeniaspirol, was added to the reaction buffer to inhibit ClpYQ via antibiotic treatment [18]. The total reaction volume was 60 µL. Digests were analyzed by SDS-PAGE, and ClpYQ activity was confirmed through hydrolysis of the fluorogenic Cbz-Gly-Gly-Leu-AMC (Millipore Sigma) substrate.

The peptide hydrolysis assay was performed as previously explained by Labana and collaborators, and the *in vitro* degradation assay was adapted from the same protocol [17]. All *in vitro* degradation assays were performed in minimum three biological replicates. Band quantification was done using ImageJ and a one-way ANOVA with a Tukey-Kramer HSD variation was used to ensure that client and non-client proteins showed a statistically significant difference in remaining protein abundance.

### Minimum inhibitory concentration

Minimum inhibitory concentration (MIC) was carried out in LB, 100 μL assays in 96-well plates. Sequential concentrations of compound (Cl-armeniaspirol or spectinomycin) were pipetted into each column through two-fold serial dilutions. 5 µL of bacterial culture already transformed with pTL12 or pTL13 (OD_600_ 0.07-0.1) was inoculated into each well and incubated at 37 °C for 16 h. Growth was observed by OD_600_ readings using a Synergy H4 microplate reader (BioTek). The MIC was determined by the lowest concentration of compound that prevented bacterial growth. All MICs were performed in three biological replicates. A summary of MICs of spectinomycin and Cl-armeniaspirol against *B. subtilis* strains transformed with expression vectors encoding DivIVA or Mbl is shown in Table S4.

### *B. subtilis* transformation

First, multimeric double-stranded DNA was generated using rolling circle amplification (RCA) since this form of DNA is easier for *B. subtilis* to uptake than monomeric double-stranded DNA [55]. Reaction mixtures were prepared with a 25 μL final volume and contained 1 X Φ29 DNA polymerase reaction buffer (NEB), 0.1 mg/mL recombinant albumin (NEB), 0.2 mM of dNTPs (NEB), 5 μM of phosphorothioate-protected random hexamers (Thermo Scientific), and 1 ng of pTL12 or pTL13 template. Reaction mixtures were primed by heating to 95 °C for 3 min and gradually cooled to 20 °C over 30 min. 5 U of Φ29 DNA polymerase was then added to the mixtures and RCA was executed at 30 °C for 18 h. Amplification was confirmed via gel electrophoresis and reactions were purified using the GFX PCR DNA and Gel Band Purification Kit (Cytiva)

To transform *B. subtilis, B. subtilis* BKE16150 (*ΔclpQ::erm*) and BKK34540 (*ΔclpP::kan*) were inoculated in 3 mL of LB supplemented with 1 µg/mL of erythromycin (for BKE16150) or 7.5 µg/mL of kanamycin (for BKK34540) overnight at 37 °C. Cultures were diluted to an OD_600_ of 0.1 in 10 mL competence media (10.7 g/L K_2_HPO_4_, 5.2 g/L KH_2_PO_4_, 20 g/L glucose, 0.88 g/L trisodium citrate dihydrate, 0.022 g/L ferric ammonium citrate, 2.5 g/L potassium aspartate, 40 mg/L L-tryptophan, 0.05% yeast extract, 10 mM MgSO_4_, 150 nM MnCl_2_) supplemented with 1 µg/mL of erythromycin (for BKE16150) or 7.5 µg/mL of kanamycin (for BKK34540) and grown to an OD_600_ of 1.5. Next, 50 ng of RCA amplified PTL12 or PLT13 was added to 120 µL of bacterial culture and incubated at 37°C (225 rpm) for 2 h. Transformation into the *ΔclpQ* mutant was plated on LB agar supplemented with 1 μg/mL erythromycin + 12.5 μg/mL lincomycin + 6 µg/mL chloramphenicol. Transformations into the *ΔclpP* mutant were plated on LB agar supplemented with 7.5 μg/mL kanamycin + 6 µg/mL chloramphenicol. Plates were incubated at 37°C overnight.

This method of transformation is similar to a previously published *B. subtilis* transformation protocol [56]. A summary of all bacterial strains used in this study is shown in Table S3.

### *In vivo* degradation assays

*ΔclpQ* and *ΔclpP* mutant cells transformed with expression vectors encoding C-terminally His-tagged Mbl or DivIVA (pTL12 or pTL13) were grown in 300 mL of LB supplemented with 6 µg/mL of chloramphenicol at 37 °C to an OD_600_ of 0.7. IPTG was then added to a final concentration of 1 mM to induce protein expression for 6 h. The MIC of spectinomycin was added after 6 h of protein expression to inhibit translation (time 0). At the same time, half the MIC of Cl-armeniaspirol was added to control cells to inhibit ClpYQ or ClpXP. At t=0, 15, and 90 min, 100 mL of cells were withdrawn, were pelleted at 4,000 rpm for 20 min, and were re-suspended in lysis buffer (100 mM Tris, 300 mM NaCl, 1 μg/mL leupeptin, 1 μg/mL pepstatin A, 1 mg/mL lysozyme, pH 8.0). Mechanical cell lysis was achieved by three 1 minute cycles of 3 s sonication and 2 s rest on ice followed by a 1 min incubation on ice. The cell debris were pelleted at 10,000 rpm for 60 min and the supernatant was mixed with 1x SDS loading buffer. Whole cell lysates were analyzed by western immunoblotting with a monoclonal anti-His antibody conjugated to HRP (Genescript).

All *in vivo* degradation assays were performed in minimum three biological replicates. Band quantification was done using ImageJ and p-values were calculated using a two-tailed Student’s t-test.

## Data availability

All data from this study are contained within the manuscript and supporting information. Any additional information required to reanalyze the data reported in this paper is available from the lead contact, Christopher N. Boddy upon request.

## Supporting information

This article contains supporting information.

## Funding and additional information

This work was funded by the Natural Sciences and Engineering Research Council of Canada and the University of Ottawa.

## Conflict of interest

The authors declare that they have no conflicts of interest with the contents of this article.

## Acknowledgements

This work was funded by the Natural Sciences and Engineering Research Council of Canada (NSERC) (RGPIN 2019-06859).

## Notes

### Competing Interest Statement

The authors have declared no competing interest.

