## supplemental information and figures for "The ATP-dependent protease ClpYQ degrades cell division proteins DivIVA and Mbl in *Bacillus subtilis*"

**Table S1. Reproduction of quantatative proteomics data for proteins described in this work** [1]. Fold-change of various proteins in the *B.subtilis* wildtype strain with Cl-armeniaspirol treatment, the *B.subtilis ΔclpP* strain, and the *B.subtilis ΔclpQ* strain relative to the the *B.subtilis* wildtype strain with no treatment is reported as the average fold change of three replicates ± standard deviation. Significance is defined as fold-change>2 and p-value<0.05 (n.d.: not detected).

| | | Cl-armeniaspirol treatment | | *ΔclpP* strain | | *ΔclpQ* strain | |
|---|---|---|---|---|---|---|---|
| **NCBI Accession** | **Description** | **Fold-change** | **p-value** | **Fold-change** | **p-value** | **Fold-change** | **p-value** |
| P80698 | Trigger factor, tig | 11.539±2.233 | 0.002 | 3.802±1.249 | 0.0001 | 0.797±1.655 | 0.349 |
| O34894 | Septation ring formation regulator EzrA, ezrA | 0.043±3.816 | 0.013 | 6.815±1.860 | 0.003 | 0.497±2.382 | 0.152 |
| P39751 | MreB-like protein Mbl, mbl | n.d. | n.d. | 16.574±2.523 | 0.005 | 14.241±1.275 | $2.197 \times 10^{-5}$ |
| P71021 | Septum site-determining protein DivIVA, divIVA | 7.360±1.285 | 0.003 | 4.584±2.176 | 0.113 | 4.792±3.146 | 0.040 |
| P80244 | ATP-dependent Clp protease proteolytic subunit ClpP, clpP | 0.893±1.949 | 0.708 | - | - | 1.261±1.268 | 0.117 |
| P39070 | ATP-dependent protease subunit ClpQ, clpQ | 0.631±2.080 | 0.211 | 0.423±2.176 | 0.057 | - | - |

**Table S2. Forward (fwd) and reverse (rev) primers used in this study.**

| **Description** | **Sequence** |
|---|---|
| *B. subtilis mbl* fwd | GTACATATGTTTGCAAGGGATATTGGTATTGACCTCG |
| *B. subtilis mbl* rev | AGGCTGAATTCCTTAGTTTGCGTTTAGGAAGCTTGTCCATATT |
| *B. subtilis divIVA* fwd | GTACATATGCCATTAACGCCAAATGATATTCACAACAAGACG |
| *B. subtilis divIVA* rev | AGGGGATCCTCCTTTTCCTCAAATACAGCGTCGACTTCATACTC |
| *B. subtilis tf* fwd | GTACATATGTCTGTAAAATGGGAAAAACAAGAAGGCAAC |
| *B. subtilis tf* rev | GTAGGATCCCGGTTTTCTACAAGAAAATCAATTGCTTTGCG |
| *B. subtilis ezrA* fwd | GTACATATGGAGTTTGTCATTGGATTATTAATTGTACTGCTTGCG |

| | |
|---|---|
| *B. subtilis ezrA* rev | AGGCTGAATTCGCGGATATGTCAGCTTTGATTTTTTCAACTGC |
| *B. subtilis clpY* fwd | AATTGCTAGCAACTTGAGGCGCATGGA |
| *B. subtilis clpY* rev | CGTAGAATTCAATATAAATTGACTTAAATC |
| *B. subtilis clpQ* fwd | CGTACATATGTCATCTTTTCATGCGACCAC |
| *B. subtilis clpQ* rev | CGTAGAATTCTCAAGCTCTTCCAGTATGATTTG |
| Cloning *divIVA* into pHT01 fwd | GGCCTGATCAATGCCATTAACGCCAAATGA |
| Cloning *divIVA* into pHT01 rev | AATTATGACGTCTTTGTTAGCAGCCGGATCT |
| Cloning *mbl* into pHT01 fwd | GGCCTGATCAATGTTTGCAAGGGATATTGG |
| Cloning *mbl* into pHT01 rev | ATTTATGACGTCTTTCGGGCTTTGTTAGCAG |

**Table S3. Bacterial strains and plasmids used in this study.**

| RESOURCE | SOURCE | IDENTIFIER |
|---|---|---|
| **Bacterial strains** | | |
| *Bacillus subtilis* 168 | Bacillus Genetic Stock Center | BGSC 1A1 |
| *Bacillus subtilis* 168 *ΔclpQ::erm trpC2* | Bacillus Genetic Stock Center | BKE16150 |
| *Bacillus subtilis* 168 *ΔclpP::kan trpC2* | Bacillus Genetic Stock Center | BKK34540 |
| **Recombinant DNA** | | |
| *mbl* in pET21b (C-term His tag, AmpR) | This study | pTL01 |
| *divIVA* in pET21b (C-term His tag, AmpR) | This study | pTL02 |
| *tf* in pET21b (C-term His tag, AmpR) | This study | pTL03 |
| *ezrA* in pET21b (C-term His tag, AmpR) | This study | pTL04 |
| *mbl* in pMGXHA (N-term His tag, AmpR) | This study | pTL05 |
| *divIVA* in pMGXHA (N-term His tag, AmpR) | This study | pTL06 |
| *tf* in pMGXHA (N-term His tag, AmpR) | This study | pTL07 |
| *ezrA* in pMGXHA (N-term His tag, AmpR) | This study | pTL08 |
| *clpQ* in pET21b (C-term His tag, AmpR) | This study | pPL29 |
| *clpY* in pET28b (N-term His tag, KanR) | This study | pPL31 |
| *mbl* in pHT01 (C-term His tag, CmR) | This study | pTL12 |
| *DivIVA* in pHT01 (C-term His tag, CmR) | This study | pTL13 |

**Table S4. MIC evaluation of spectinomycin and Cl-armeniaspirol against *B. subtilis* strains transformed with expression vectors encoding DivIVA or Mbl.**

| | Minimum Inhibitory Concentration (μg/mL) | |
|---|---|---|
| **Strain** | **Cl-armeniaspirol** | **Spectinomycin** |
| *ΔclpP*[A] | 4 | 64 |
| *ΔclpP*[B] | 2 | 32 |
| *ΔclpQ*[A] | 2 | 128 |
| *ΔclpQ*[B] | 2 | 32 |

[A]Transformed with expression vector encoding DivIVA

[B]Transformed with expression vector encoding Mbl

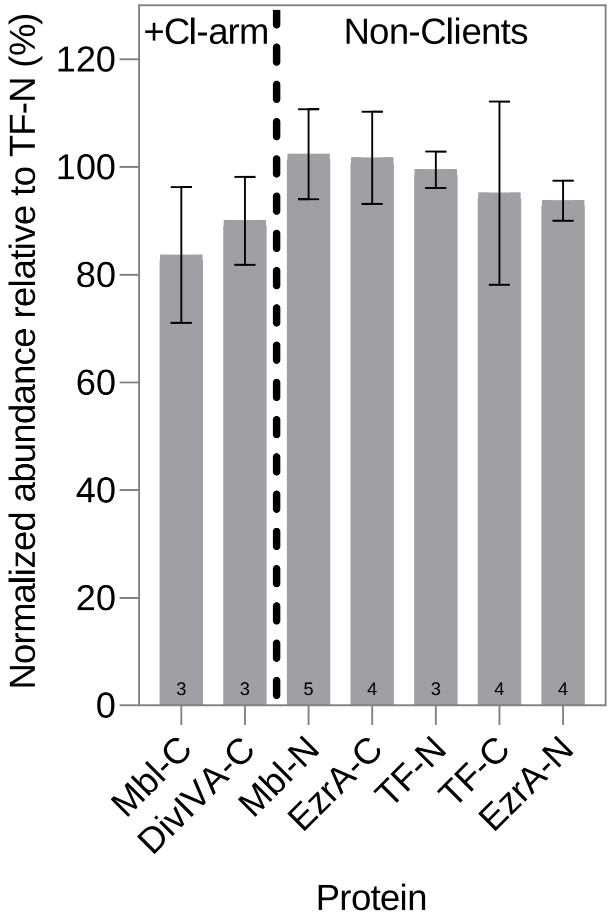


**Figure S1. Inhibition of *in vitro* ClpYQ-mediated degradation of C-terminally His-tagged Mbl and DivIVA by Cl-armeniaspirol (Cl-arm).** Recombinant purified C-terminally His-tagged Mbl and DivIVA were incubated with recombinant purified ClpYQ and 4X the $K_I$ of Cl-armeniaspirol for ClpYQ (25 µM of Cl-armeniaspirol) for 24 h at 37°C. Protein abundance is compared to non-client proteins. The terminus onto which the His tag was fused is indicated by the C or N suffix on the protein names. Digests were analyzed by SDS-PAGE. Each band was normalized to a loading control before quantifying the amount of protein remaining in the reaction mixture relative to a no-ATP control. Protein abundance was then calculated as a percentage relative to the TF-N negative control. Error bars represent the standard deviations from n=3-5 independent replicates, and all groups were compared to each other using a one-way ANOVA with a Tukey-Kramer HSD variation to ensure no statistically significant difference in remaining protein abundance between client proteins in the presence of Cl-armeniaspirol and non-client proteins but a statistically significant difference between client proteins in the absence and presence of the antibiotic. No degradation of client proteins was observed in the presence of Cl-armeniaspirol.

**Table S5. Results of Tukey-Kramer HSD variation run on one-way ANOVA data.** Statistically significant for $\alpha \leq 0.05$.

| Substrate 1 | Substrate 2 | p-value | Statistical Significance |
|---|---|---|---|
| DivIVA-C | Mbl-C | 0.0037 | S |
| DivIVA-C | TF-C | 0.11 | NS |
| DivIVA-C | EzrA-C | 0.012 | S |
| DivIVA-C | TF-N | 0.044 | S |
| DivIVA-C | EzrA-N | 0.37 | NS |
| DivIVA-C | Mbl-N | 0.020 | S |
| Mbl-C | TF-C | 9.99E-06 | S |
| Mbl-C | EzrA-C | 8.83E-07 | S |
| Mbl-C | TF-N | 2.31E-06 | S |
| Mbl-C | EzrA-N | 5.6E-05 | S |
| Mbl-C | Mbl-N | 1.46E-06 | S |
| TF-C | EzrA-C | 0.98 | NS |
| TF-C | TF-N | 1.0 | NS |
| TF-C | EzrA-N | 1.0 | NS |
| TF-C | Mbl-N | 0.99 | NS |
| EzrA-C | TF-N | 0.99 | NS |
| EzrA-C | EzrA-N | 0.76 | NS |
| EzrA-C | Mbl-N | 1.0 | NS |
| TF-N | EzrA-N | 0.97 | NS |
| TF-N | Mbl-N | 1.0 | NS |
| EzrA-N | Mbl-N | 0.86 | NS |

S: Significant
NS: Not Significant

**Table S6. Amount of Mbl and DivIVA degraded in the *ΔclpP B.subtilis* strain after 90 minutes.**

| | % Degradation | |
|---|---|---|
| **Substrate** | **-Cl-armeniaspirol** | **+Cl-armeniaspirol** |
| DivIVA | 4 | 64 |
| Mbl | 2 | 32 |
